# The effects of resource subsidy duration on detritus-based stream ecosystem: a stream mesocosm experiment

**DOI:** 10.1101/770370

**Authors:** Takuya Sato, Rui Ueda, Gaku Takimoto

## Abstract

Most of the resource subsidies are temporally variable, and studies have revealed that ecological processes can be mediated by the temporal attributes of subsidies, such as timing and frequency. Less studies have, however, examined the effects of the subsidy duration, an another major temporal attribute, on consumer populations, communities and ecosystem functions. Using an outdoor mesocosm experiment, we demonstrated that, even with the same total amounts, the prolonged subsidy let large-stage fish effectively monopolize the subsidy over small-stage fish, while the pulsed subsidy allowed small-stage fish to increase the ingestion rate of the subsidy. This effect resulted in causing weaker indirect positive effects on *in-situ* benthic prey and a leaf breakdown rate with the prolonged subsidy than with the pulsed-subsidy although it depended on dominant benthic prey species having different edibility. Increasing evidences have shown that global warming would not only advance, but also prolong the growing seasons, which may, in turn, make subsidies more prolonged. The ecological significance of the subsidy duration might be common in nature, and should be incorporated to better understand ecological processes in spatially and temporally coupled ecosystems.

## Introduction

Resource subsidies that link heterogeneous habitats are ubiquitous in natural ecosystems (Polis et al. 1997) and often vary temporally, such as returning salmons (Gende et al. 2002), emergences of arthropods (Nakano and Murakami 2001) and storm-driven sea weeds (Spiller et al. 2010). Teasing apart the effects of temporal attributes (magnitude, timing and duration) of the resource subsidies is thus important to understand population-, community- and ecosystem-dynamics (Yang et al. 2008; Richardson and Sato 2015), as well as to forecast the effects of climate changes on those dynamics (Larsen et al. 2016; O’Gorman 2016). Previous studies have demonstrated that seasonal timing of resource subsidies is a critical component in structuring consumer populations and seasonal organization of communities (Takimoto et al. 2002; Leroux and Loreau 2013; Sato et al. 2016). On the other hand, seasonal duration may also be important because it would regulate consumer responses if, for example, pulsed inputs satiate consumer’s capacity of handling and/or assimilating subsidies (Armstrong and Bond 2013; Uno 2016), which may in turn affect communities and ecosystem processes (Holt 2008). While global warming is well-known to advance peak timing of growing season, it also leads longer growing seasons (CaraDonna et al. 2014). This suggests that resource subsidies may be prolonged under climate changes. However, until recently less studies have directly tested the effects of subsidy duration on the dynamics of consumer populations and their communities (Richardson and Sato 2015).

A key mechanism by which the subsidy duration affects consumer responses would be contrasting resource partitioning among consumer individuals between pulsed and prolonged subsidies even with the same total amounts of subsidies. Specifically, pulsed subsidies would result in an equivalent resource partitioning among consumers because individuals cannot fully monopolize the subsidies due to their limited handling and/or assimilation capacities (Uno 2016). Contrary to the pulsed subsidies, prolonged subsidies would give an opportunity for some individuals to gain more subsidies over other individuals due to intraspecific variations in handling/ assimilation capacities (Uno 2016) and ability of interference competition (Sato and Watanabe 2014). As a consequence, amount of body size variation of consumers may become higher with the prolonged subsidies than with the pulsed subsidies.

The consumer responses to the subsidy duration may further affect communities and ecosystem functions in recipient ecosystems. If the prolonged subsidies let competitively superior individuals to dominate the subsidies, inferior individuals would maintain their predation pressure on *in-situ* prey (i.e., size-dependent functional response: Sato and Watanabe 2014). Consequently, the positive indirect effects of the subsidies on *in-situ* prey are likely to be weaker with the prolonged subsidies than with the pulsed subsidies.

Here, we provide the first experimental test of the effects of the subsidy duration on stream ecosystems by using an outdoor mesocosm experiment, in which we directly manipulated the subsidy duration (pulsed vs. prolonged) of terrestrial invertebrate input into the mesocosms. In this study, we tested (1) whether the variation of individual growth rate and amount of body size variation of consumers (red-spotted masu salmon *Oncorhynchus masou ishikawae*) become higher with the prolonged subsidy than with the pulsed subsidy; if so, (2) cascading indirect effects of the subsidy on *in-situ* prey (benthic invertebrates) and an ecosystem function (leaf break-down rate) are weaker with the prolonged subsidy than with the pulsed subsidy (Sato and Watanabe 2014).

## Materials and Methods

### Experimental design of resource subsidy duration

To test the duration effect of a seasonally occurring terrestrial invertebrate subsidies on stream ecosystems, we conducted an outdoor mesocosm experiment, in which we artificially added commercially available mealworms (larvae of beetle *Tenebrio molitor*) into the mesocosms at different duration (pulsed = 30 days vs. prolonged = 90 days), keeping the total amounts and peak timing of the mealworm inputs. Specifically, in the pulsed-subsidy mesocosms, mealworms were added intensively but briefly at the rate of 90 mg dry mass/ m^2^/ day for 30 days at a middle of the experimental period (i.e., from 30th to 60th days). On the other hand, mealworms were added throughout the experimental period (i.e., 90 days) at the rate of one third of the pulsed treatment (i.e., 30 mg/m^2^/day) in the prolonged-subsidy mesocosms. To separate the duration effects from the effects of the resource subsidy per se, we used control mesocosms where no mealworms were added. The durations (30 and 90 days) and input rates (30 and 90 mg/m^2^/day) are set within the natural ranges of the terrestrial invertebrate inputs in temperate streams (Nakano and Murakami 2001; Kawaguchi et al. 2002; Sato et al. 2011; Sato et al. 2016). See Sato et al. 2016 for rational and more detailed methods about the mealworm addition.

### Experimental preparations

The mesocosm experiment was conducted from 20 August to 18 November, 2016 in two stream sides of the nearby 2^nd^-order streams (hereinafter, st. Azodani and st. Mizukoshi) in the Wakayama Research Forest Station (WRFS) of Kyoto University (34° 3’53.4”N, 135° 31’28.4”W). The six commercially available pools (L × W × H: 300 × 200 × 70 cm) were set at each streamside (12 pools in total). Each pool was fed by stream water (40 cm in water depth) that was siphoned from the upstream of the experimental sites in each stream with approximately 0.6 m^3^/ sec discharge, which make mealworms naturally drift within the pool. At least one month before the onset of the experiment, we added natural gravels into each pool at several centimeters depth, which allowed natural colonization of benthic invertebrates (i.e., *in-situ* prey). In addition to the natural colonization, we transferred benthic invertebrates collected from 6.0 m^2^ of natural stream beds in a nearby stream by randomly collecting 67 quadrats (surber sampler: 0.09 m^2^ in area with 250 μm in mesh size). This would make the community composition of benthic invertebrate community similar among mesocosms at the onset of the experiment. The four blocks (L × W × H: 39 × 10 × 19 cm) were evenly placed on the gravel beds to make the habitat complexity that could relax the interference interaction among fish, otherwise the interaction would be strong due to the artifactitious simple habitat structure within the pools.

One week before the onset of the experiment, we captured red-spotted masu salmon *Oncorhynchus masou ishikawae* (hereinafter, masu salmon) in a nearby stream and released them into each pool. Four small fish (average ± SE: 90 ± 5 mm in fork length, n = 12 mesocosms) and two large fish (121 ± 5 mm) were released into each pool to promote the development of natural social hierarchies (Sato and Watanabe 2014). This population structure (2:1 of small and large fish) and density (1.0 fish/ m^2^) were within the range of the natural population in the study site (2–4 small and 1–2 large fishes per pool and 0-2.3 fish/ m^2^ in pool habitats). These masu salmons were anaesthetized, measured (fork length to the nearest 1 mm and weighed to the nearest 0.1 g), and individually marked by inserting a visible implant elastomer (VIE) into dorsal fin before release in each pool.

### Consumer responses to the subsidies

To evaluate the contribution of each prey category (mealworms, other terrestrial invertebrates and benthic invertebrates) to fish diets, masu salmon were captured by electrofishing once during the last week of each subsidy treatments (i.e., ≈60 days and 90 days in the pulsed- and prolonged-subsidy mesocosms, respectively). Although a single assay of fish diet would inevitably be a snapshot, we did not repeat the assay during each subsidy period because we were afraid that multiple-electrofishing over such a short period in a small mesocosm would adversely affect masu salmon’s behavior and/or physiology. To compensate for potential bias caused by the snapshot assay, we carefully chose weather conditions (i.e., fair weather) and a time period (14:00 to 16:00) for catching masu salmon. We collected the stomach contents of 4-6 masu salmon per each mesocosm (>20 masu salmon per treatment). After capture, masu salmon were measured as described above, and their stomach contents were quickly pumped. The stomach samples were preserved in 70% ethanol and identified to order to distinguish their sources (mealworms, terrestrial or benthic). Relative growth rate for individual masu salmon (66 fish in total) was calculated by the following equation (Marczak and Richardson 2008): Growth rate = ((m_2_/m_1_)^1/t^ − 1) × 100, where m_1_ and m_2_ are the individual masses of masu salmon at the beginning and the end of the experiment, and *t* is the duration of the experiment (i.e., no. of days = 90).

### Responses of benthic community and an ecosystem function

To measure the responses of benthic invertebrates (mainly, shredder invertebrates) and a leaf break-down rate, we placed three litter packs (made of ≈ five air-dried leaves of *Acer rufinerve*, a common riparian tree, with stems tied together, and approximately 0.73 ± 0.10 g in weight) in each pool at the onset of the experiment. Each litter pack was directly anchored to a cobble so that the litter packs were located on the gravel beds throughout the experiment. At the end of the experiment, individual litter pack was collected delicately by a small hand-net (0.5 mm in mesh size). The litter packs were then placed in a tray and well-rinsed to remove all invertebrates from the litter packs. All benthic invertebrates were preserved in 70% ethanol, and individuals were identified to the family, genus or species depending on the taxa. The litter packs were dried for 72 h at 30 °C and weighed in the laboratory. We did not measure ash-free dry mass (AFDM) because litter packs were well-rinsed in the field and very little inorganic material was expected. Leaf-break down rate (*L*_*R*_) was then calculated by comparing changes in mass (Benfield 2006).

### Data Analysis

All statistical analysis were performed using R version 3.4.3. (R Core Team 2017), with alpha of 0.05. We used generalized linear mixed models (GLMM; lme function in the R package “nlme”), with site (st. Azodani and st. Mizukoshi) and blocks (n = 2) within each site as random effects, to test whether the ingestion rates of mealworms and benthic invertebrates, and specific growth rate of individual masu salmon was respectively affected by treatment (pulsed vs. prolonged vs. control), fish stage (large vs. small stages) and their interaction. The interaction term was excluded from the final model if it does not significantly maximize the log-likelihood of the model in the log-likelihood ratio test (Crawley 2007). Normal error distributions and identity link functions were used. To meet the assumption of the model, we log-transformed the continuous response variable if necessary (i.e., ingestion rates of mealworms and benthic invertebrates) and confirmed the asymptotic normality of error by a Q-Q plot. When the treatment effect was significant, it could be that only one treatment was different from the others. Therefore, we tested if combining some treatments further maximized the log-likelihood of the GLMM model by using the log-likelihood ratio test in lme. Specifically, we compared the full model with three combined models (‘pulse’ + ‘prolonged’ vs. control; ‘pulse’ + ‘control’ vs. prolonged; or ‘prolonged’ + ‘control’ vs. pulse).

The coefficient of variation (CV) for fork length of masu salmon in each mesocosm was calculated to evaluate if the amount of body size variation of masu salmon become different among treatments toward the end of the experiment. To further evaluate the population-level consequences of the subsidy durations, we pooled the data for four replicates in each treatment and tested if the frequency distribution of the body size differed significantly among treatments.

For community/ ecosystem responses, our primary objective was to simply test the effects of subsidy duration by assuming random variations nested to the site and block. However, a slight difference of light condition between the two sites (openness: st. Azodani = 40%; st. Mizukoshi = 60%) inevitably differentiated dominant benthic invertebrates (see results in details) in the two sites, which could result in affecting the effects of the functional response of masu salmon to the subsidy. Therefore, we compared a GLMM (glmer function in the R package “glmmML”) simply testing the treatment effects by considering site and block as random factors with an alternative model that included treatment and site as fixed factors and block within each site as a random factor by using log-likelihood ratio test. The optimal model in this comparison was shown in the results.

## Results

### Consumer responses to pulsed vs. prolonged subsidies

When the mealworm subsidy was available, consumption rate of the mealworms was on average 2.9 times higher for masu salmon in the pulsed-subsidy mesocosms [mean ± SE (n = 4 mesocosms): 0.15 ± 0.06 mg / 100 mg dry mass of fish] than in the prolonged subsidy mesocosms (0.05 ± 0.01; GLMM, treatment: *F*_*1, 38*_ = 20.32, *P* < 0.0001), as well as 4.1 times higher in the large-fish stage than in the small-fish stage (large: 0.26 ± 0.16; small: 0.07 ± 0.05; *F*_*1, 38*_ = 29.80, *P* < 0.0001; Table 1a). However, the effects of the subsidy duration and fish stage were additive, not synergetic (the interaction term was not significantly improved the model fit: log-likelihood ratio test: *P* = 0.32; Fig 1A), suggesting that the large-fish stage did not further increase the ingestion rate of the increased subsidy in the pulsed-subsidy mesocosms compared with prolonged-subsidy mesocosms. As a consequence, the small-stage fish in the pulsed-subsidy mesocosms consumed, on average, 3.2 times more mealworms (0.10 ± 0.05 mg/ 100 mg mass of fish) than those in the prolonged subsidy mesocosms (0.03 ± 0.01) (Fig. 1A), although fish could gain mealworms for three-times longer period in the prolonged subsidy than in the pulsed subsidy mesocosms.

**Table 1.**
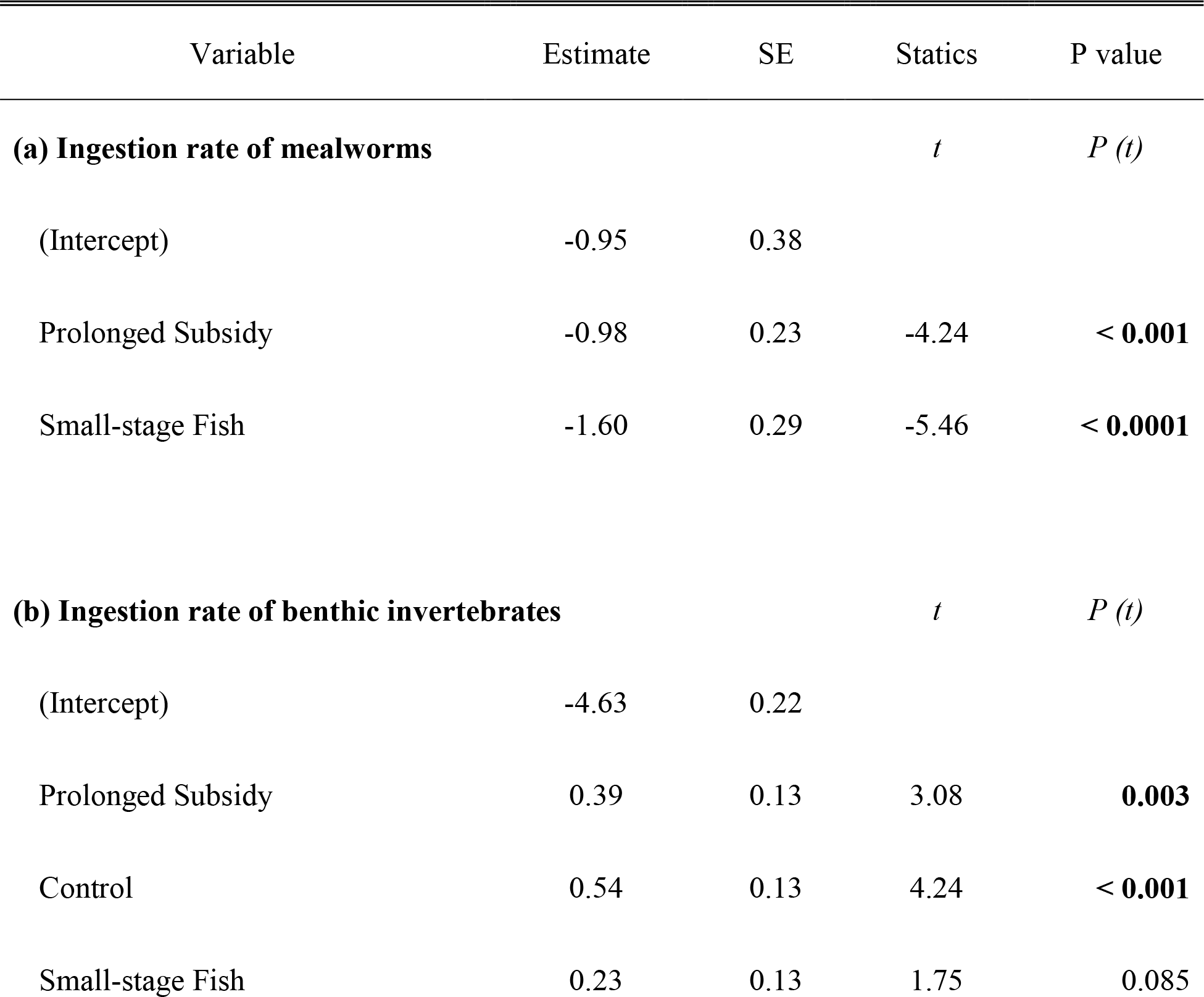
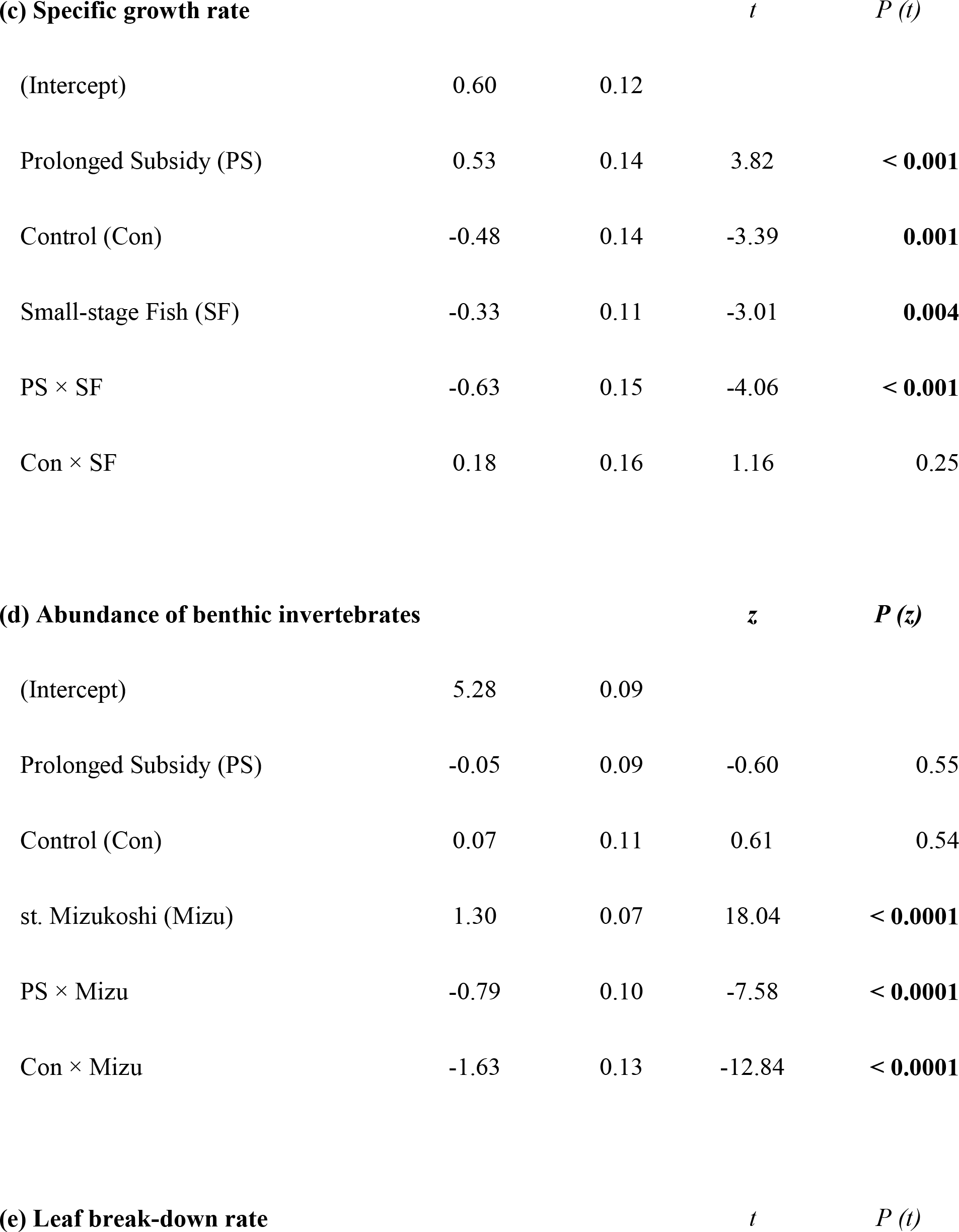
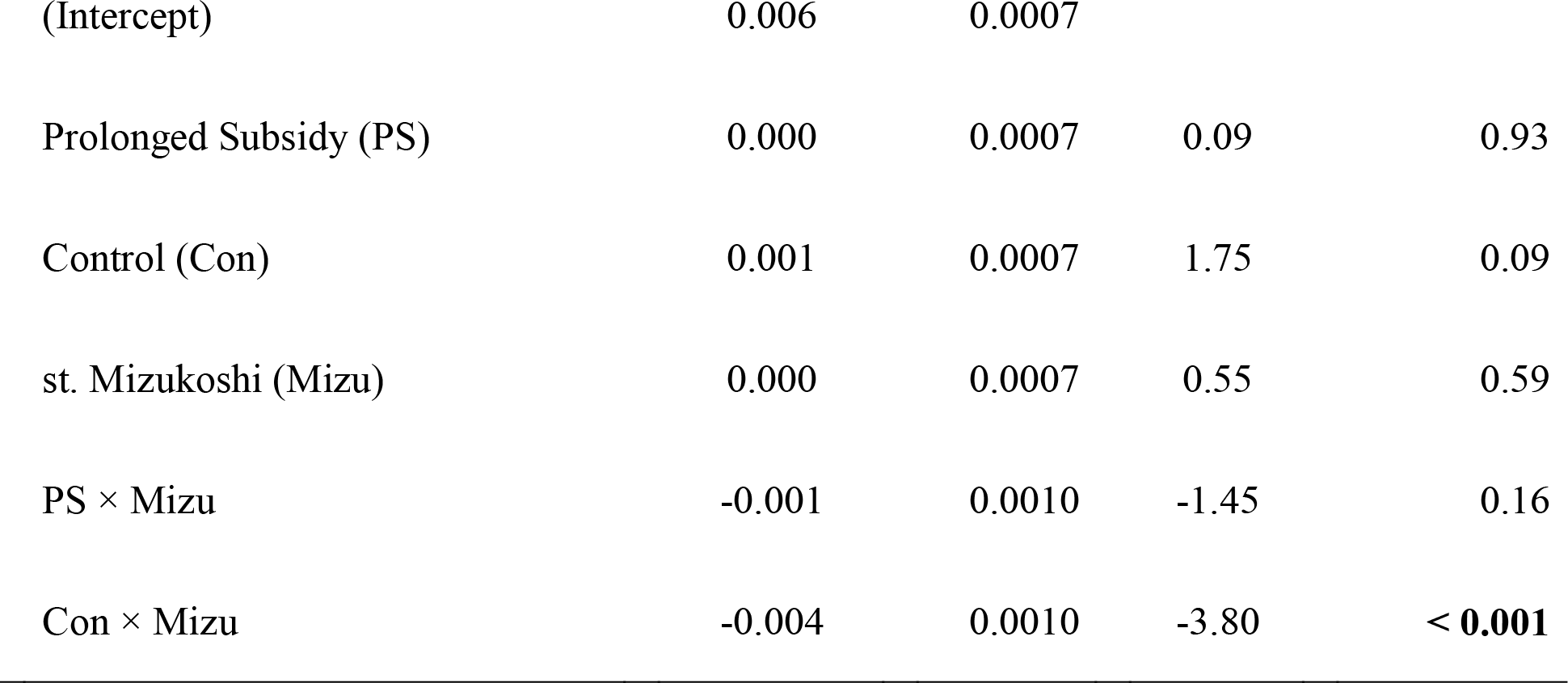
Results of GLMM models for ingestion rates of mealworms (a) and benthic invertebrates (b), specifice growth rate of individual masu salmon (c), abundance of benthic invertebrates in a litter pack (d), and leaf break-down rate (e). In the GLMM analyses, Pulsed subsidy, Large-stage fish and st. Azodani were used as the contrasts with the subsidy treatments, fish stage and site, respectively.

**Figure 1.**
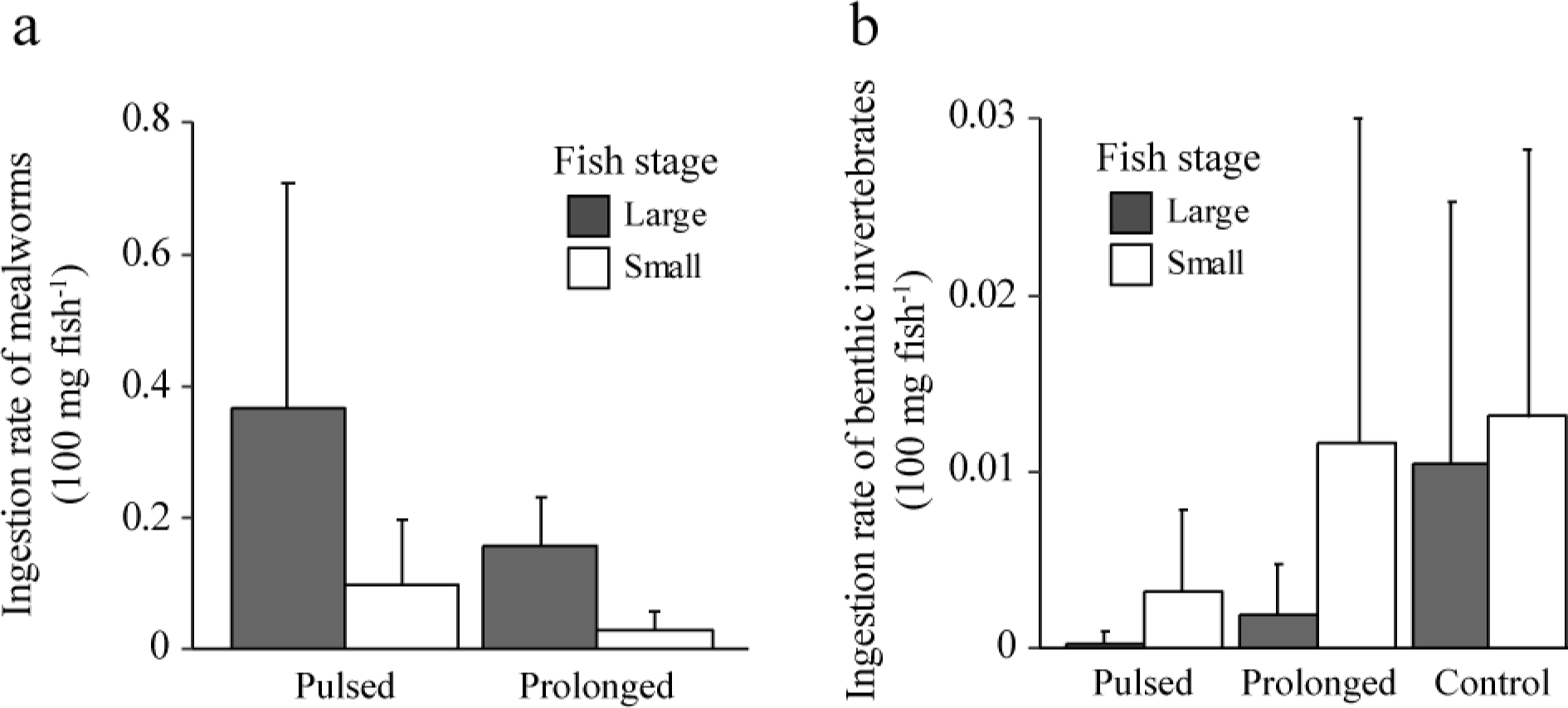
The effects of treatments on the ingestion rates (mean ± SE, n = 4) of mealworms (a) and benthic invertebrates (b) by large and small masu salmons, respectively.

Growth responses of masu salmon mirrored the way of the resource partitioning among masu salmon under different subsidy durations. Specifically, either subsidies increased the growth rate of masu salmon in comparison with control mesocosms (GLMM, treatment: *F*_*2, 59*_ = 20.94, *P* < 0.0001; Fig. 2A). However, the large-stage salmon that received the prolonged subsidy grew, on average, 89% faster than those that received the pulsed subsidy (treatment × fish stage: *F*_*2, 59*_ = 15.17, *P* < 0.0001; Fig. 2A, Table 1c). On the other hand, small-stage salmon that received the prolonged subsidy grew, on average, 33% slower than the small salmon that received the pulsed subsidy (Fig. 2A).

**Figure 2.**
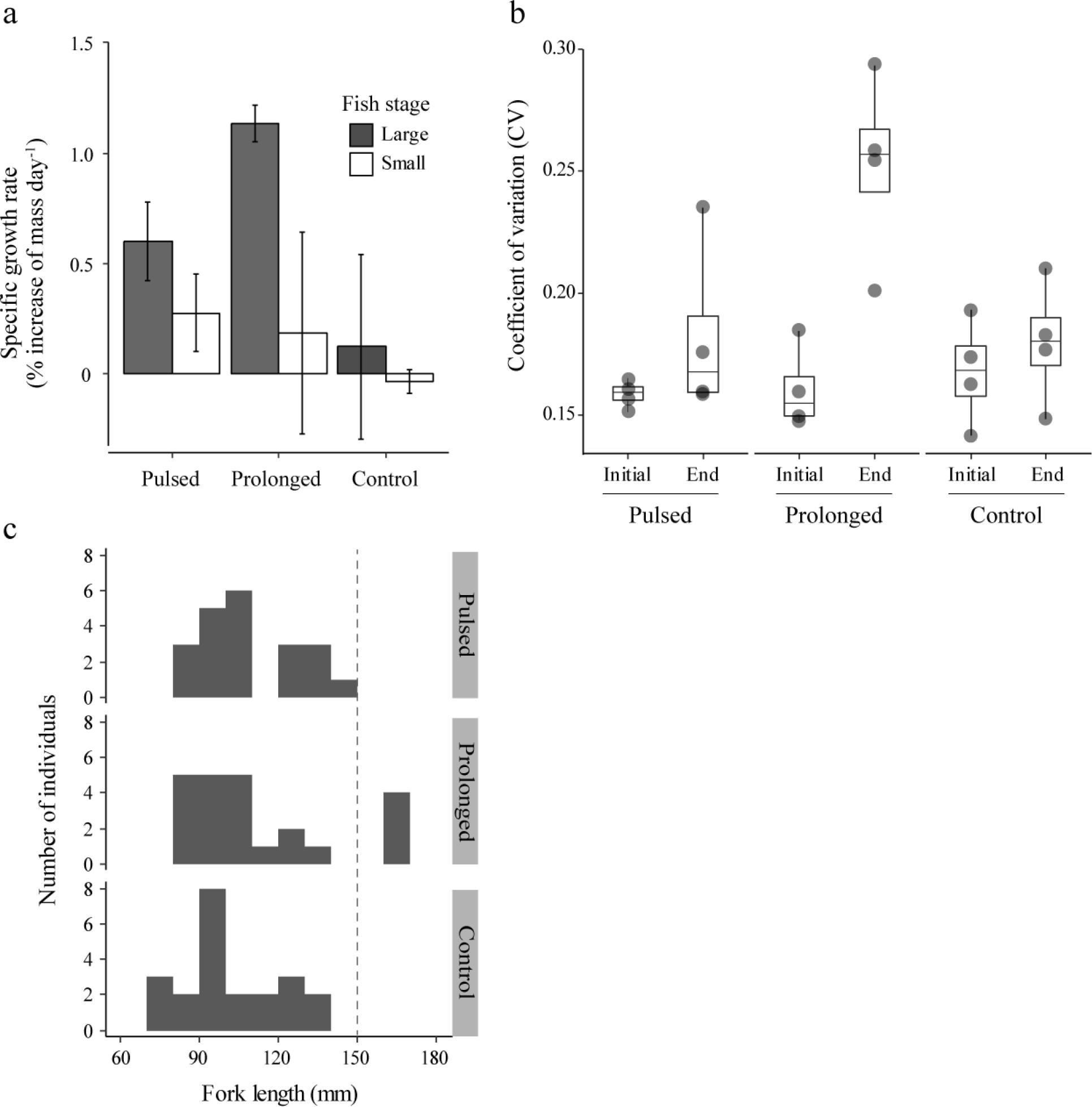
The effects of treatments on individual growth rate (mean ± SE, n = 4) (a), coefficient of variation for body size (b) and distribution of body size (c) of masu salmon. In the panel (b), points represent the CVs in each mesocosm (n = 4). In the panel (c), a vertical dashed line shows the average body size of mature males and females in the studied population (Sato et al. unpublished data).

A population-level consequence of receiving different subsidy durations was found for the amount of body size variation of masu salmon (Fig. 2B, C). For instance, the body size of masu salmon was more variable in the prolonged-subsidy mesocosms (average CV = 0.25, n= 4) than those observed in the pulsed- and control-mesocosms (average CVs: pulsed = 0.18; control = 0.18; Fig. 2B). In the prolonged-subsidy mesocosms, most of the large-stage fish reached sexually mature size (≈150 mm in FL in the studied population; Ueda et al. unpublished data) within the three months, while no large-stage fish did in the pulsed- and control-mesocosms (Fig. 1C).

### Cascading effects on in-situ prey and leaf break-down rate

Overall, there was a significant negative correlation between the ingestion rates of mealworms and benthic invertebrates, i.e., *in-situ* prey (Pearson’s correlation: *r* = −0.31, *P* = 0.047, n = 42), suggesting the functional response of masu salmon to the mealworm subsidy. The ingestion rates of benthic invertebrates were significantly lower in either subsidy mesocosms than those in the control mesocosms [mean ± SE (n = 4 mesocosms); 0.0118 ± 0.007 mg/ 100 mg dry mass of fish; GLMM, treatment: *F*_*2, 59*_ = 9.71, *P* = 0.0002; Table 1b). The trend was more pronounced in the pulsed-subsidy mesocosms (0.0017 ± 0.002) compared to the prolonged-subsidy mesocosms (0.0068 ± 0.008) because clustering the two subsidy treatments significantly reduced the model fit; log-likelihood ratio test: *P* < 0.05). Ingestion rate of benthic invertebrates highly varied among individuals and thus was not significantly related to the fish-stage (*F*_*1, 59*_ = 3.08, *P* = 0.09; Table 1b), although smaller individuals tended to ingest more benthic invertebrates (correlation between body size and benthic ingestion rate: *r* = −0.24, *P* = 0.052, n = 42).

In the litter packs, midge larvae (Chironomidae species) that colonized the litter surface occupied a large proportion (97%) of the total abundance of benthic invertebrates in the st. Mizukoshi. On the other hand, isopods (*Asellus* sp.) that prefer the inside of litter pack dominated the benthic invertebrates (96 %) in the st. Azodani. Probably due to this difference in microhabitat use between the two dominant taxa, a possible indirect positive effects of the subsidies on the total abundance of benthic invertebrate was found in the st. Mizukoshi, but not in the st. Azodani (Fig. 3A, Table1d). In st. Mizukoshi, total abundances of benthic invertebrates were higher in either subsidy mesocosms than those in the control mesocosms (Fig. 3A, Table 1d). Furthermore, the benthic invertebrates were, on average, 31% lower in the prolonged-subsidy mesocosms than those in the pulsed-subsidy mesocosms, indicating indirect positive effect in the prolonged-subsidy than in the pulsed-subsidy mesocosms (Fig 3A).

**Figure 3.**
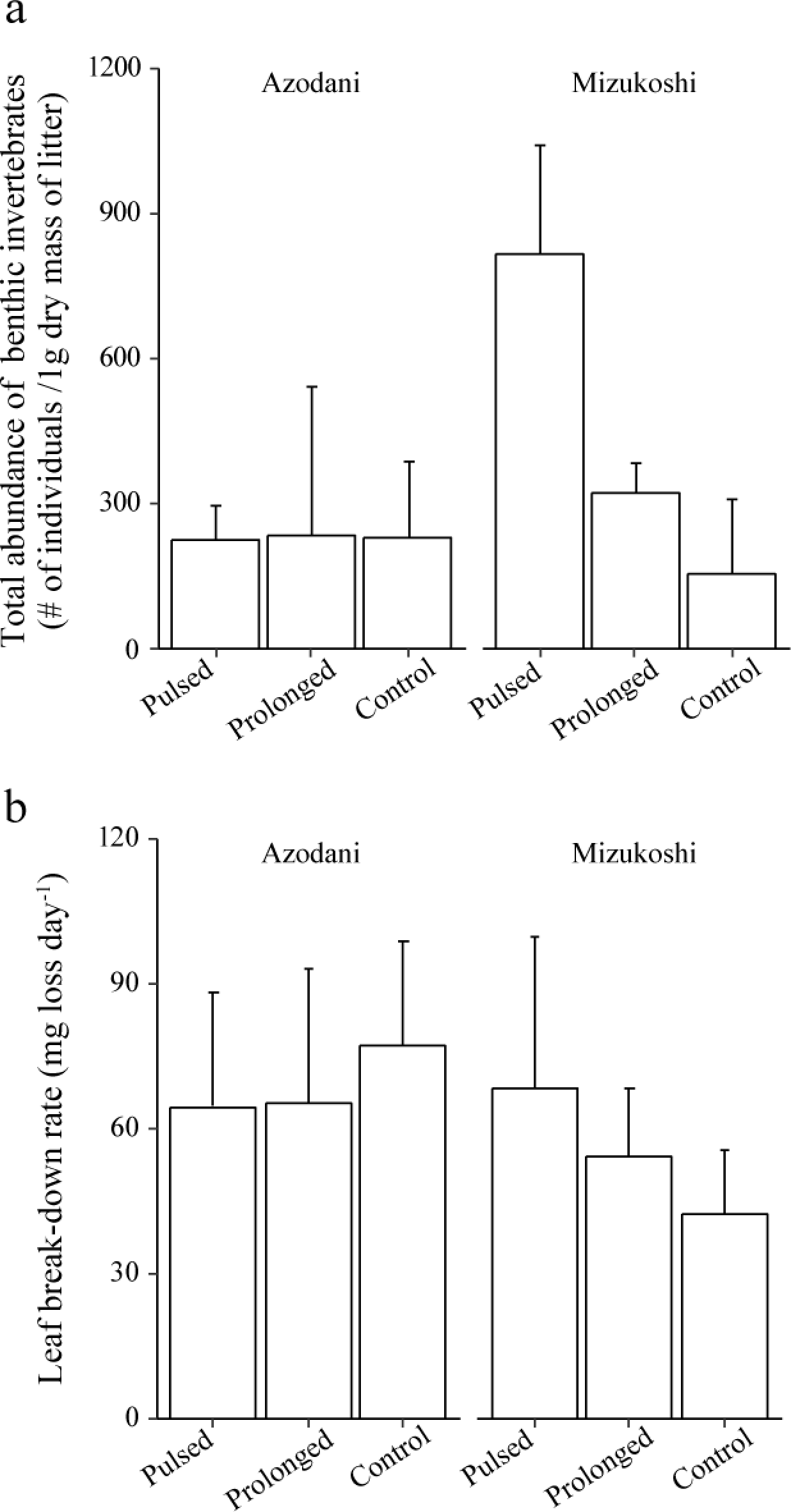
The effects of treatments on total abundances of benthic invertebrates (a) and leaf break-down rate (b) in the st. Azodani and st. Mizukoshi.

Further cascading effect on leaf breakdown rate was also found in the st. Mizukoshi, but not in the st. Azodani (GLMM: treatment × site, *F*_*2, 29*_ = 7.33, *P* = 0.003; Fig. 3B, Table 1e). In the st. Mizukoshi, the leaf breakdown rate was significantly faster in either subsidy mesocosms than those in the control mesocosms (Fig. 3B, Table 1e). However, the leaf breakdown rate was, on average, 20% slower in the prolonged-subsidy than in the pulsed-subsidy mesocosms, suggesting the cascading effect in the former than in the latter subsidy mesocosms (Fig 3B; Table 1e).

## Discussion

Most of the resource subsidies that link heterogeneous habitats are temporally variable (Yang et al. 2008; Richardson and Sato 2015). Studies have begun to elucidate how ecological processes can be mediated by the temporal attributes of subsidies, such as timing and frequency (Marczak and Richardson 2008; Yee et al. 2012; Sato et al. 2016). However, until recently, less studies have directly tested the effects of the subsidy duration, an another major temporal attribute, on consumer populations, communities and ecosystem functions (but see Uno 2016 for consumer individuals). Using an outdoor mesocosm experiment, we demonstrated that, even with the same total amount, the prolonged terrestrial invertebrate subsidy generated more size-varied salmonid population and conditionally caused weaker indirect positive effects on *in-situ* benthic prey and a leaf breakdown rate compared with the pulsed-subsidy. To the best of our knowledge, this is the first study thoroughly testing the effects of subsidy duration on consumer population, community and ecosystem function.

### Duration-dependent resource partitioning and population structure of consumer

Resource partitioning can strongly regulate individual growth rate and size-structure of consumer populations, which in turn affect community organizations and stability (Bolnick et al. 2011; Rudolf and Lafferty 2011). In riparian ecosystems, the terrestrial invertebrate is commonly a dominant prey resource for stream fishes from summer to autumn seasons (Nakano and Murakami 2001) when our experiment was conducted. In the mesocosm experiment, we found that the prolonged mealworm subsidy let large-stage fish effectively monopolize the subsidy over small-stage fish, while the pulsed subsidy allowed small-stage fish to increase the ingestion rate of the subsidy. The way of these resource partitioning under different subsidy durations were well-translated to the individual growth rate of fish consumers. Consequently, the prolonged subsidy generated more size-varied masu salmon population than the pulsed subsidy did. Masu salmon reached mature body size only in the prolonged subsidy mesocosms, suggesting that the prolonged subsidy may facilitate the numerical response of masu salmon.

While we did not conduct the detailed behavioral and physiological assays, the size-based interference competition, as well as the food saturation and assimilation capacity of large-stage fish, would cause the contrasting resource partitioning under the two different subsidy durations. Specifically, the size-based interference competition is known to strongly regulate resource partitioning in stream salmonid fishes, including red-spotted masu salmon (Nakano 1995; Nakano et al. 1999). In this regard, we previously demonstrated that large masu salmon less foraged during daytime with high subsidy input, which resulted in an increased ingestion rate of subsidy for small masu salmon due to the weakened interference from large to small fish (Sato and Watanabe 2014). Furthermore, although masu salmon experienced the same total amount of subsidy, the specific growth rate and the eventual body size of large-stage fish were smaller in the pulsed-subsidy than those in the prolonged-subsidy mesocosms, suggesting that an assimilation capacity constrained the growth rate of the large-stage fish during the pulsed subsidy.

The interference competition is a ubiquitous form of competition among animals especially at higher trophic levels. Assimilation capacity is a common constrain for organisms to capitalize the pulse of food availability in the feast and famine environment (Armstrong and Schindler 2011; Armstrong and Bond 2013). Together with the potential variation in subsidy durations in nature (Yang et al. 2008; Richardson and Sato 2015), the results we observed would be general and important in understanding a manner of population regulation in spatially coupled ecosystems.

### Short-term effects on community and ecosystem function

The contrasting resource partitioning under different subsidy durations differently cascaded down to affect the abundance of benthic invertebrates (i.e., *in-situ* prey) and leaf breakdown rate in mesocosms in the st. Mizukoshi, but not in the st. Azodani. While the interacting effects of treatment and site may seem complicated, this could be explained by the differences of the functional traits (i.e., habitat preference and mobility) of the dominant benthic invertebrates in the two sites. Specifically, immobile midge larvae that predominantly colonized the litter surfaces must suffer from the fish predation in the st. Mizukoshi. As a consequence, the functional response of masu salmon would elicit the positive indirect effects of the subsidy on the benthic invertebrates and hence the leaf breakdown rate in the st. Mizukoshi. Furthermore, the prolonged subsidy let masu salmon (especially smaller salmon) ingest more benthic invertebrates than those in the pulsed subsidy, which resulted in causing a weaker positive indirect effects in the prolonged-subsidy compared with the pulsed-subsidy mesocosms. Those processes were consistent with the processes demonstrated in our previous study, in which the stage-specific functional responses of masu salmon mediated the strength of trophic cascade (Sato and Watanabe 2014).

On the other hand, nocturnal *Asellus* sp. usually colonized inside of the litter packs and would be invulnerable to the fish predation. In this case, the functional response of masu salmon, if any, would not be efficiently transferred to mediate the indirect effects. Such the predator-specific vulnerability of primary consumers has been known to mediate the strength of trophic cascade both in a grazing-based and a detritus-based stream food webs (Ruetz et al. 2002; Power et al. 2008). We could not explain what environmental factors derived the dominances of different invertebrate taxa in the two sites. In future, the effects of the subsidy duration on trophic cascade would be more mechanistically elucidated by considering functional traits of benthic invertebrates and environmental gradients that would regulate them (McGill et al. 2006 TREE).

### Implications for the long-term effects on community and ecosystem function

While we empirically tested the indirect effects of the subsidy duration on recipient stream ecosystems (i.e., benthic prey and leaf-beak down rate), it would be practically difficult to extrapolate their long-term consequences solely by the short-term experiment. The prolonged subsidy weakened the short-term positive indirect effects by dampening the functional response of fish consumers (especially small fish) compared with the pulsed subsidy. These short-term positive indirect effects may be canceled out over time due to consumers’ numerical response. For instance, the prolonged subsidy let competitively superior individuals (i.e., large-fish stage) grow more and reach mature-size faster in our experiment, which can cause stronger numerical response of consumer with the prolonged subsidy than with the pulsed subsidy in our theoretical model (Takimoto & Sato XXX). On the other hand, if the growth rate of small-fish stage is weakened with the prolonged subsidy than with the pulsed subsidy (partially supported in our experiment), the prolonged subsidy may result in weaker numerical response compared with the pulsed subsidy. Taken together, the long-term effects of subsidy duration would be expressed as a combination of how subsidy duration modifies the short-term functional responses and long-term numerical responses of consumers to subsidy inputs (Takimoto & Sato XXX; Piovia-Scott et al. 2019).

### Conclusions

Spatial community ecology has recently made progress towards understanding the community dynamics that receive temporal resource subsidies by considering relative timing between subsidies and *in-situ* resources (Takimoto et al. 2002; Leroux and Loreau 2012), timescales of subsidies relative to those of the consumers’ numerical responses (Takimoto et al. 2009) and timing-dependent consumer responses (Wright et al. 2013; Sato et al. 2016). In addition to those studies, our results demonstrated that the duration of subsidy can also be a critical component determining the consumer’s population structure, dynamics of communities and ecosystem functions in recipient ecosystems. Phenological events (e.g., salmon spawning run and arthropod emergences) of individuals that generate temporal subsidies can be synchronized (pulsed) or desynchronized (prolonged) depending on the degree of spatial/ temporal environmental heterogeneities (Uno and Power 2015; Uno 2016; Armstrong et al. 2016; Deacy et al. 2017). Furthermore, global warming would lead longer growing seasons (CaraDonna et al. 2014), which may, in turn, make subsidies more prolonged. Therefore, the results we observed might be widespread in nature and should be considered in predicting ecosystem processes in seasonal environments. By integrating key ecological processes acting within and across seasons into a theoretical framework, one can develop a general framework to better predict community organizations and their interactions with ecosystem functions in spatially and temporally coupled ecosystems.

## ACKNOWLEDGEMENT

We thank S. Naman for edits and comments that improved the manuscript. Ryo Arai, Yoshikazu Asano, Atsushi Hasegawa, Tomonori Katsuyama, Hisaya Uenichi and Shin Ukaji are acknowledged for their assistance in setting up and running the experiment in Wakayama Forest Research Station, FSERC, Kyoto University. This work was supported by JSPS KAKENHI Grant Number JP 15H04422.

## LITERATURE CITED

Armstrong, J. B. and M. H. Bond. 2013. Phenotype flexibility in wild fish: Dolly Varden regulate assimilative capacity to capitalize on annual pulsed subsidies. Journal of Animal Ecology 82:966–975.

Armstrong, J. B. and D. E. Schindler. 2011. Excess digestive capacity in predators reflects a life of feast and famine. Nature 476:84–87.

Armstrong, J. B., G. Takimoto, D. E. Schindler, M. M. Hayes, and M. J. Kauffman. 2016. Resource waves: phenological diversity enhances foraging opportunities for mobile consumers. Ecology 97:1099–1112.

Benfield, E. 2006. Decomposition of leaf material. Pages 711–720 in F. R. Hauer and G. A. Lamberti, editors. Methods in stream ecology. Elsevier Inc., Burlington, MA.

Bolnick, D. I., P. Amarasekare, M. S. Araújo, R. Bürger, J. M. Levine, M. Novak, V. H. Rudolf, S. J. Schreiber, M. C. Urban, and D. A. Vasseur. 2011. Why intraspecific trait variation matters in community ecology. Trends in Ecology and Evolution 26:183–192.

CaraDonna, P. J., A. M. Iler, and D. W. Inouye. 2014. Shifts in flowering phenology reshape a subalpine plant community. Proceedings of the National Academy of Sciences 111:4916–4921.

Crawley, M. J. 2007. The R book. John Wiley and Sons, UK.

Deacy, W. W., J. B. Armstrong, W. B. Leacock, C. T. Robbins, D. D. Gustine, E. J. Ward, J. A. Erlenbach, and J. A. Stanford. 2017. Phenological synchronization disrupts trophic interactions between Kodiak brown bears and salmon. Proceedings of the National Academy of Sciences 114:10432–10437.

Gende, S. M., R. T. Edwards, M. F. Willson, and M. S. Wipfli. 2002. Pacific Salmon in Aquatic and Terrestrial Ecosystems: Pacific salmon subsidize freshwater and terrestrial ecosystems through several pathways, which generates unique management and conservation issues but also provides valuable research opportunities. Bioscience 52:917–928.

Holt, R. D. 2008. Theoretical perspectives on resource pulses. Ecology 89:671–681.

Kawaguchi, Y. and S. Nakano. 2001. Contribution of terrestrial invertebrates to the annual resource budget for salmonids in forest and grassland reaches of a headwater stream. Freshwater Biology 46:303–316.

Larsen, S., J. D. Muehlbauer, and E. Marti. 2016. Resource subsidies between stream and terrestrial ecosystems under global change. Global Change Biology 22:2489–2504.

Leroux, S. J. and M. Loreau. 2012. Dynamics of Reciprocal Pulsed Subsidies in Local and Meta-Ecosystems. Ecosystems:1–12.

Marczak, L. B. and J. S. Richardson. 2008. Growth and development rates in a riparian spider are altered by asynchrony between the timing and amount of a resource subsidy. Oecologia 156:249–258.

McGill, B. J., B. J. Enquist, E. Weiher, and M. Westoby. 2006. Rebuilding community ecology from functional traits. Trends in Ecology and Evolution 21:178–185.

Nakano, S. 1995. Competitive interactions for foraging microhabitats in a size-structured interspecific dominance hierarchy of two sympatric stream salmonids in a natural habitat. Canadian Journal of Zoology 73:1845–1854.

Nakano, S., K. D. Fausch, and S. Kitano. 1999. Flexible niche partitioning via a foraging mode shift: a proposed mechanism for coexistence in stream-dwelling charrs. Journal of Animal Ecology 68:1079–1092.

Nakano, S. and M. Murakami. 2001. Reciprocal subsidies: dynamic interdependence between terrestrial and aquatic food webs. Proceedings of the National Academy of Sciences 98:166–170.

O’Gorman, E. J. 2016. It’s only a matter of time: the altered role of subsidies in a warming world. Journal of Animal Ecology 85:1133–1135.

Piovia-Scott, J., L. H. Yang, A. N. Wright, D. A. Spiller, and T. W. Schoener. 2019. Pulsed seaweed subsidies drive sequential shifts in the effects of lizard predators on island food webs. Ecology Letters (https://doi.org/10.1111/ele.13377).

Polis, G. A., W. B. Anderson, and R. D. Holt. 1997. Toward an integration of landscape and food web ecology: the dynamics of spatially subsidized food webs. Annual Review of Ecology and Systematics:289–316.

Power, M. E., M. S. Parker, and W. E. Dietrich. 2008. Seasonal reassembly of a river food web: floods, droughts, and impacts of fish. Ecological Monographs 78:263–282.

R Core Team. 2017. R: A Language and Environment for Statistical Computing. R foundation for statistical computing, Vienna, Austria. URL http://www.R-project.org/.

Richardson, J. S. and T. Sato. 2015. Resource subsidy flows across freshwater–terrestrial boundaries and influence on processes linking adjacent ecosystems. Ecohydrology:In press.

Rudolf, V. H. and K. D. Lafferty. 2011. Stage structure alters how complexity affects stability of ecological networks. Ecol Lett 14:75–79.

Ruetz, C. R., R. M. Newman, and B. Vondracek. 2002. Top-down control in a detritus-based food web: fish, shredders, and leaf breakdown. Oecologia 132:307–315.

Sato, T., R. W. Elsabaawi, K. Campbell, T. Ohta, and J. S. Richardson. 2016. A test of the effects of timing of a pulsed resource subsidy on stream ecosystems. Journal of Animal Ecology 85:1136–1146.

Sato, T. and K. Watanabe. 2014. Do stage-specific functional responses of consumers dampen the effects of subsidies on trophic cascades in streams? Journal of Animal Ecology 83:907–915.

Sato, T., K. Watanabe, M. Kanaiwa, Y. Niizuma, Y. Harada, and K. D. Lafferty. 2011. Nematomorph parasites drive energy flow through a riparian ecosystem. Ecology 92:201–207.

Spiller, D. A., J. Piovia-Scott, A. N. Wright, L. H. Yang, G. Takimoto, T. W. Schoener, and T. Iwata. 2010. Marine subsidies have multiple effects on coastal food webs. Ecology 91:1424–1434.

Takimoto, G., T. Iwata, and M. Murakami. 2002. Seasonal subsidy stabilizes food web dynamics: balance in a heterogeneous landscape. Ecological Research 17:433–439.

Takimoto, G., T. Iwata, and M. Murakami. 2009. Timescale hierarchy determines the indirect effects of fluctuating subsidy inputs on in situ resources. The American Naturalist 173:200–211.

Uno, H. 2016. Stream thermal heterogeneity prolongs aquatic-terrestrial subsidy and enhances riparian spider growth. Ecology 97:2547–2553.

Uno, H. and M. E. Power. 2015. Mainstem-tributary linkages by mayfly migration help sustain salmonids in a warming river network. Ecology Letters 18:1012–1020.

Yang, L. H., J. L. Bastow, K. O. Spence, and A. N. Wright. 2008. What can we learn from resource pulses. Ecology 89:621–634.

Yee, D. A. and S. A. Juliano. 2012. Concurrent effects of resource pulse amount, type, and frequency on community and population properties of consumers in detritus-based systems. Oecologia 169:511–522.

